# Integrating Behavioural Science and Epidemiology to Improve Early Detection of Zoonotic Swine Influenza in the Netherlands

**DOI:** 10.1101/2025.04.03.646521

**Authors:** Juliette Fraser, Ewa Pacholewicz, Peter Hobbelen, Thomas Hagenaars, Ron Bergevoet, Michel Counotte

## Abstract

**Background and Objectives:** The Netherlands faces zoonotic disease risks due to its dense human and livestock populations. The 2009 H1N1 outbreak highlighted the pandemic potential of influenza virus reassortment. Effective preparedness requires integrating behavioural and epidemiological models. Human behaviour, shaped by personal, social, and institutional factors, is critical in detecting, intervening, and treating diseases. Using the Theory of Planned Behaviour (TPB), a framework was developed integrating knowledge from the TPB to improve early detection and response, using (zoonotic) swine influenza as a case study.

**Material and Methods:** Within the framework we defined the desired outcome: timely detection and notification of symptomatic (and hypothetical zoonotic) swine influenza to prevent its spread. Actions, such as symptom recognition and disease reporting, were linked to key drivers extracted from the TPB and disease transmission modelling. Expert elicitation estimated the likelihood of action for different farmer profiles, while disease transmission modelling assessed farm-to-farm spread probabilities. Simulations integrated these probabilities to evaluate intervention effectiveness across different scenarios.

**Results:** The framework successfully combined behavioural science and epidemiology, offering nuanced estimates of intervention effectiveness. For early detection, 95% of farmers were estimated to notify their veterinarian within 13 days post-infection. Key factors influencing action included symptom recognition and disease spread extent. The farmer profiles influenced response likelihood, while human infections linked to outbreaks had minimal impact. Farm density and assumptions about transmission probabilities significantly affected the likelihood of spread before notification.

**Discussion and Conclusion:** The framework provides a systematic approach for integrating social and epidemiological insights to support evidence-based policies. The work can be further enhanced by complementing expert judgement with more extensive stakeholder surveys, randomized scenario presentations, and immersive methods. This pragmatic tool aids policymakers in designing targeted interventions for zoonotic disease preparedness.

## Introduction

In the last decade, the Netherlands experienced outbreaks and the circulation of (zoonotic) pathogens in livestock, for example, Q-fever [1], SARS-CoV2 [2], avian influenza [3–5] and swine influenza [6]. The latter highlights the pandemic risk posed by the emergence of zoonotic strains, as demonstrated by the 2009 H1N1/09-influenza outbreak in humans [7]. The widespread endemic circulation of influenza A viruses in pigs, combined with the probability of reassortment with human or avian influenza, could lead to the emergence of an outbreak similar to the 2009 H1N1 outbreak, posing significant public health risks. The Netherlands, characterized by its high population density of both humans and livestock, faces a heightened risk of zoonotic disease emergence [7]. To prevent the transmission of diseases from animals to humans, it is essential to maintain vigilance and implement well-coordinated actions in response to outbreak signals. To support the policy makers in the preparedness and response phases of an outbreak infectious disease modelling is commonly applied both in the public and veterinary health domains [8, 9]. Understanding disease transmission dynamics, predicting the course of outbreaks, and predicting or evaluating the effect of interventions are important aims to inform policy and guidelines. To achieve these aims, mathematical modelling is often employed as a key tool allowing to quantify the transmission dynamics from field and/or experimental data, and subsequently extrapolate to new situations for which direct observational data is absent. In such model extrapolations, often the dynamics of the spread of the disease are simulated, using scenario assumptions for (changes in) the characteristics of the at-risk population, environment and/or intervention measures.

In addition, in preparedness for, and response to infectious disease outbreaks, human behaviour plays a crucial role. Human action is needed to detect symptoms, to implement interventions, and to seek treatment, and thus shapes the course of an outbreak. For example, individuals must seek diagnosis or treatment, or report disease presence through various means. The decision to act and the behaviours displayed are influenced by multiple drivers at different levels: personal characteristics, social environment, and the broader context encompassing value chain partners, organizations, institutions, and governmental bodies. Interventions aimed at increasing pandemic preparedness will thus almost always require the inclusion of incentives that promote desired human behaviour, including compliance with advice and regulations [10]. In the context of zoonotic diseases farmers are on the front lines of outbreaks and their actions might impact the initial diagnosis and (prevent) the spread of the disease. Understanding famers’ responses and thus key leverage points for interventions will increase pandemic preparedness and enable better anticipation of interventions’ effectiveness upon outbreak.

So far many of the outbreaks mentioned above of contagious livestock diseases have been well studied from either an epidemiological or behavioural perspective, but seldomly using integrated approaches. We currently lack approaches in which behavioural models are employed in interaction with infectious disease models in a generalizable way [11]. Such incorporation is crucial and ensures that human behaviours related to infection prevention are appropriately represented, thereby enhancing the models’ accuracy and predictive power [11]. The challenges of incorporation of behavioural aspects into epidemiological models, amongst other include exploration: 1) how and to what extent behaviour should be modelled, and 2) the level of detail required to model differences in behaviour [12]. For this exploration, quantitative parameters of behaviour could be identified with the help of behavioural theories that provide insights into human behaviour through frameworks that conceptualize individual decision-making [11]. For example, the Theory of Planned Behaviour (TPB) [13] offers a structured framework to understand the determinants of human actions. and has been widely applied to study various behaviours of farmers [14] throughout addressing three main constructs: attitude towards the behaviour, subjective norms and perceived behavioural control [13]. Developed in the 1980s by Icek Ajzen, the Theory of Planned Behaviour (TPB) is based on the principle that intention is the primary determinant of behaviour. This intention is influenced by three key factors: attitude, subjective norms, and perceived behavioural control [13]. In the context of zoonotic disease prevention, the TPB can clarify how farmers’ attitudes towards measures, the social pressures they experience, and their perceived control over implementing these measures influence their decision-making related to the threat of livestock infectious diseases.

Here, we explore integration of concepts from behavioural sciences (TPB) and epidemiology (disease transmission modelling). We propose a framework bridging both disciplines and providing both inventory as well as quantification of drivers that trigger early detection of zoonotic disease outbreaks based on swine influenza as a case study.

## Methods

We outline a generic approach to evaluate the effectiveness of key actions in achieving desired outcomes within the scope of pandemic preparedness, where we integrate concepts from behavioural science and infectious disease transmission modelling. This methodology is grounded in challenges identified in the literature [12] and insights gathered from a collaborative workshop involving experts from both the social sciences and the fields of infectious diseases and epidemiology. The approach comprises the steps depicted in Figure 1.

**Figure 1.**
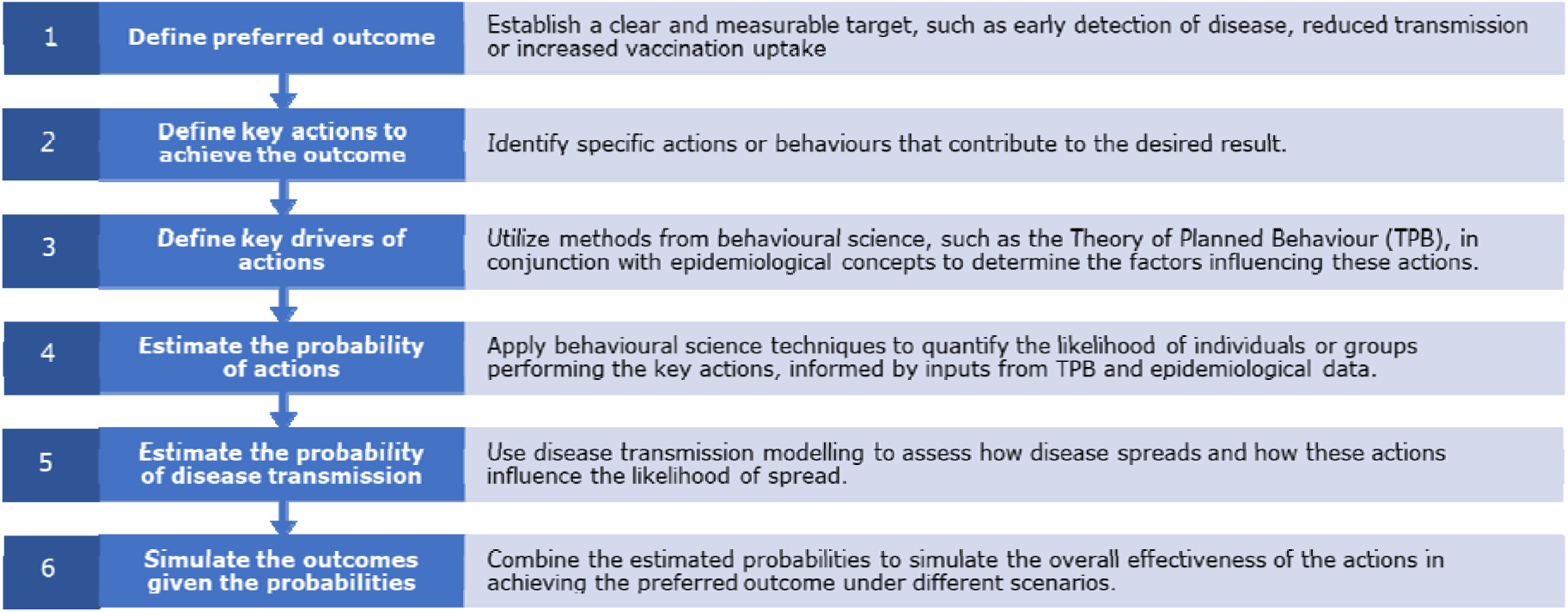
Generic approach to integrating concepts from behavioural science and infectious disease transmission modelling for outbreak and pandemic preparedness and response.

This structured framework integrates insights from behavioural science and epidemiology with quantitative modelling to systematically evaluate the potential impact of interventions on pandemic preparedness. Variability and uncertainty in the estimation of probabilities (points 4 and 5) is addressed through scenario modelling, allowing to account for the influence of different predictors and variables on these probabilities.

Throughout this work, we used ‘zoonotic swine influenza’ as a case-study to illustrate concepts and to have a concrete example for stakeholders to interact with to quantify behavioural intention. Since currently circulating subtypes of swine influenza (H1N1, H3N2 and H1N2) are seldomly causing human cases [15], we thus refer here to ‘zoonotic swine influenza’ as a hypothetical strain with increased pig-human transmissibility, for the sake of providing an example that is close to reality and facilitates increasing our pandemic preparedness.

### Definition of the preferred outcome and key actions to achieve the outcome

We defined successful ‘early detection of zoonotic swine influenza’ as the detection of a zoonotic swine influenza outbreak on a farm before it was able to transmit to other farms. We further defined this as a chain of actions that needed to be taken by different stakeholders before the veterinary authority would have received the signal that ‘zoonotic swine influenza’ was present on this farm. The actions are conditional on each other, meaning that subsequent action will only be taken once a previous action was taken. Table 1 provides an overview of the actions and the stakeholders that need to take the action in other achieve the end-goal: Early detection.

**Table 1.**
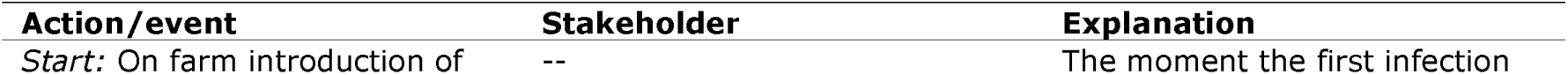
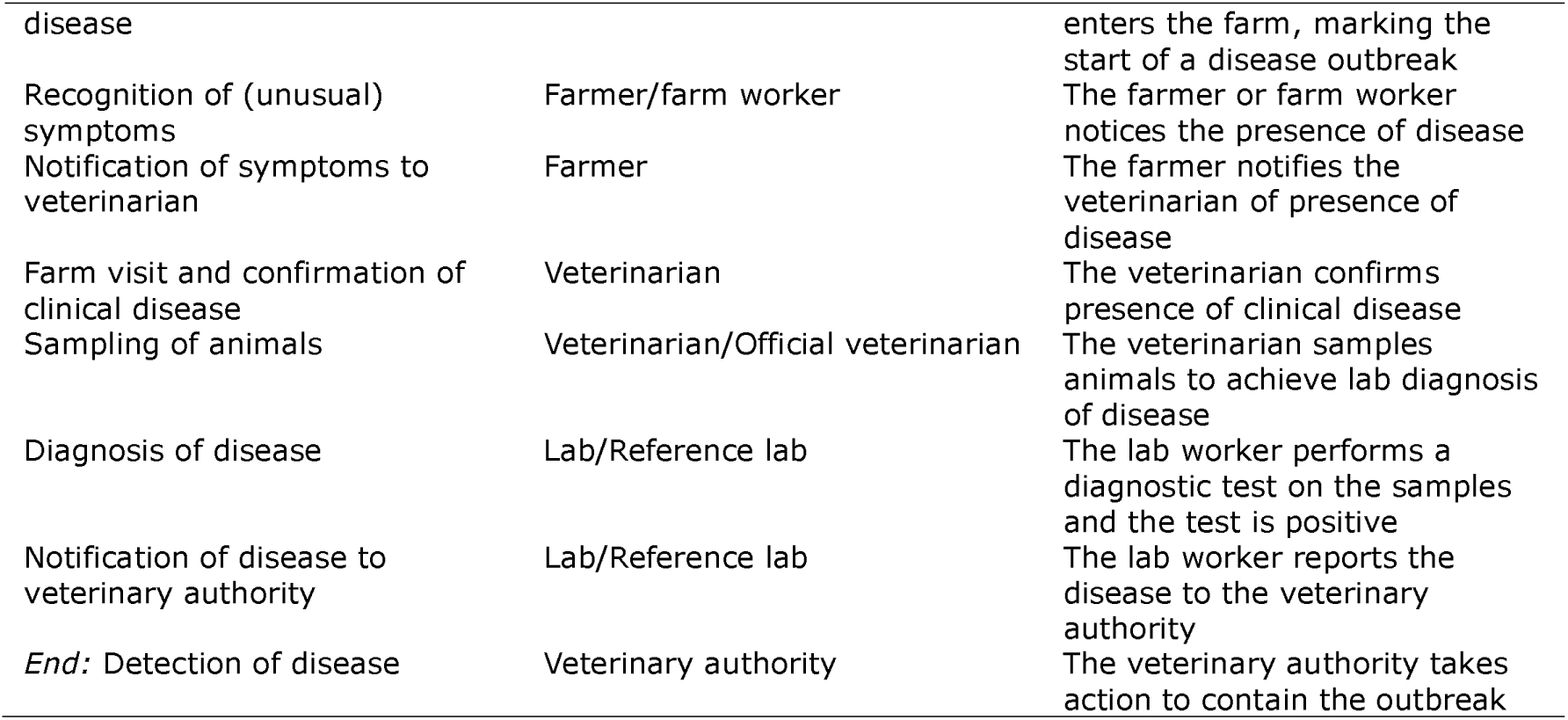
Overview of actions and stakeholders to achieve early detection of zoonotic swine influenza on a farm.

To illustrate the framework’s application, we focused on two primary actions of the farmer/farm worker: ‘Recognition of symptoms’, and ‘Notification of their veterinarian’. These behaviours were selected as they represent the initial and most direct steps leading to the detection of a disease outbreak on a farm.

### Definition of key drivers of actions

The drivers of actions serve as the fundamental components of human behaviour that influence farmers’ decision-making processes during a potential disease outbreak and can impact whether an action will happen or not. By identifying and categorizing these drivers, we aim to create a comprehensive model that captures the complex interplay of factors affecting farmers’ behaviours.

To categorize the key drivers of the actions, we applied the TPB: Intention is the primary determinant of following a behaviour, which is influenced by the constructs of attitude, subjective norms and perceived behavioural control [13]: Attitude pertains to an individual’s perception on whether a particular behaviour contributes positively or negatively to their life. Subjective norms describe the influence of social pressures manifest and the extent to which the opinion of others impact an individual’s decisions. Finally, perceived behavioural control refers to an individual’s sense of control over their ability to achieve goals and perform behaviours that depend on a series of interconnected actions [16].

The foundation of the theory includes external variables and background factors such as demographic details (gender, age, education, occupation, socioeconomic status, religion, etc.), personality traits (openness, conscientiousness, agreeableness, extraversion, neuroticism, etc.), and environmental influences like the physical environment and access to resources. These elements impact all other parts of the behavioural model, acting as underlying influences on the decision maker’s environment [16].

To make the TPB constructs more applicable, we incorporated the TPB constructs into three distinct farmer profiles. These profiles represent typical patterns of attitudes, subjective norms and perceived behavioural control that could be observed among farmers. They provide a concrete way to understand how different farmers might respond to a disease outbreak based on their individual characteristics and perceptions. Each profile embodies a unique combination of attitudes towards farm management practices, influence of subjective norms, and perceived ability to implement control measures and adapt to challenges.

We developed the following three profiles based on literature and expert input:

*1)* The *‘Family-oriented farmer’* is characterized by their deep-rooted commitment to their local community and environment. These farmers exhibit a strong sense of stewardship and prioritize the stability of their farms over expansion, often with significant family involvement [17–20].
*2)* The *’Business-oriented farmer’* is driven by profit and entrepreneurship. They are integrated into global market chains and focus on expanding and scaling up their operations [17–19, 21].
*3)* The ‘*Farmer without successor’* – was developed to provide a more comprehensive understanding of the diverse approaches to farming. In this profile, the farmer aims to cut costs and boost output, often sacrificing long-term viability for immediate financial gains. They are less likely to be influenced by their peers’ opinions, and they feel overwhelmed by farming challenges and lack confidence in decision-making due to limited resources and support. This profile highlights the varied strategies farmers might adopt in response to contemporary challenges such as financial pressures and resource constraints, which are not fully captured by the two other profiles and allowing for a more nuanced analysis of farmer behaviour.

These profiles are stylized representations and individual farmers may exhibit characteristics of both profiles. Their specific traits and motivations can vary widely depending on personal circumstances, local context, and the type of farming practiced.

We hypothesized that additional and specific factors related to disease spread could influence human behaviour, primarily through their impact on risk perception. These factors include the on-farm disease progression quantified through the number of infected animals on a farm, the occurrence of human infections (potentially) associated with the outbreak, and the availability of knowledge from previous similar outbreaks.

Risk perception, while not a core element of the TPB, is assumed to shape a farmer’s attitude towards disease control measures [22]. It can be defined as the subjective evaluation of a potential threat by an individual, based on the perceived likelihood of a threat occurring and the perceived impact if it does occur. In farm management, risk perception shapes a farmer’s beliefs about the consequences of risk-management practices and the values attached to those consequences, thereby influencing their attitude towards implementing these practices [23]. The addition of context-specific risk perception drivers enables to explore how outbreak characteristics influence behaviour through their impact on risk perception. Table 2 gives an overview of the drivers that we further define below to estimate the probability of action. In this table, we distinguish between TPB constructs (represented by the farmer profiles) and specific risk perception drivers. This distinction allows for a more nuanced analysis of how outbreak characteristics influence behaviour, while still maintaining the theoretical framework of the TPB.

**Table 2.** Drivers that can influence the probability of action.

| Driver | Scenarios | Levels |
| --- | --- | --- |
| Attitude | Three farmer profiles | 'Family-oriented farmer', |
| Subjective norms |  | 'Business-oriented farmer', |
| Perceived behavioural control |  | 'Farmer without successor' (3) |
| Risk: On-farm disease progression | Epidemic progression on the farm | Day 10, 15, 20 (3) |
| Risk: Human infections | Human cases linked to farm | True/false (2) |
| Risk: Context | Increased local threat | True/false (2) |

### Estimation of probability of action: Behavioural intention

#### Survey instrument

In order to test our methodology and to estimate the probability of action (noticing symptoms and reporting disease to the veterinarian), we performed an expert elicitation. We opted for expert elicitation and not for a large stakeholder survey, since our main objective was to illustrate the conceptual framework.

#### Measurement

To measure the relationship between drivers and the probability of action, we presented combinations of scenarios to stakeholders (Table 2). We developed a survey instrument in which stakeholders could provide estimates on the probability that a farmer would take action given certain circumstances. Per scenario, we provided a description of the farmers personality profile, the progression of disease, and contextual details. The progression of disease as number of pigs with typical influenza symptoms on the farm was presented as a table in which for day 10, 15 and 20 after the introduction of the infection, the total number of infected animals were provided. These were derived from a within-farm disease transmission model (described in detail below). Presence or absence of human infections related to the outbreak were displayed in the same table as a separate column. The contextual information whether an earlier outbreak of zoonotic swine influenza did occur was conveyed as well (See Supplement S2 for more details).

Stakeholders were asked to provide an estimate of a probability of the two farmer actions (recognition of symptoms and reporting these to the veterinarian) at three different time-points during a hypothetical outbreak. The profiles were presented to the expert in a concise three-paragraph format, with each paragraph linked to one of the three TPB constructs. This format illustrates whether the hypothetical farmer demonstrates a high or low level of the associated construct. The information was conveyed in a presentation in which profiles were represented by a pictogram and text. During the survey, the experts were presented the pictograms (Figure 2).

**Figure 2.**
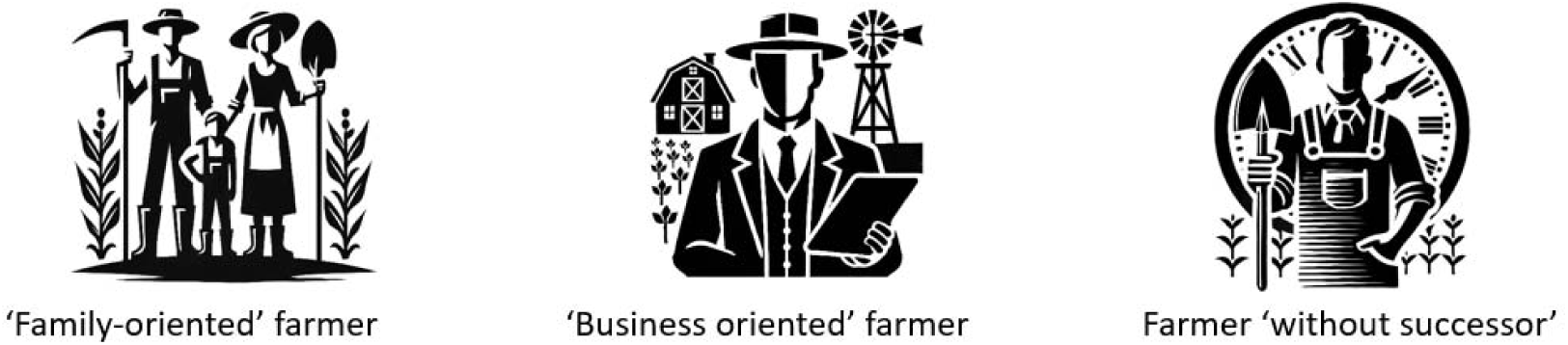
Graphical depiction of the different farmer profiles.

Additionally, we asked the stakeholders several validation questions in which they were asked to predict the response of the different farmer profiles. These questions were targeted to investigate the understanding of the respondents of the levels of subjective norms, attitude, and perceived behaviour control of the farmer profiles. The survey responses were analysed based on a theoretical hypothesis that each farmer profile would display distinct behavioural traits aligned with the TPB. Specifically, it was hypothesised that openness to technology adoption (attitude), influence of peers’ opinion (subjective norms), and confidence in farming methods (perceived behavioural control) would vary consistently across profiles, reflecting common stereotypes or expectations associated with each type. These theoretical foundations guided the development and scoring of the validation questions to assess alignment with these anticipated behavioural patterns.

Each response to the validation question was scored on a Likert scale, ranging from ’strongly disagree’ to ’strongly agree’, to capture the degree of alignment with each statement. The scale enables a nuanced assessment of respondents’ perceptions, with scores reflecting varying levels of agreement or disagreement. These scores provide a quantifiable means to assess the extent to which respondents perceived each profile in line with the theoretical expectations associated with subjective norms, attitudes toward innovation, and perceived behavioural control. We conducted the survey with three experts from the Netherlands in the field of swine health. See Supplement S2 for the full survey instrument.

### Analysis

We quantified the relationship between the logit of the estimated probability of action and the drivers, by assuming a linear relationship, using beta-regression [24]. We used a logit link function to map the linear predictor ni to the interval (0, 1). The relationship between the predictor and the variables is described in EQ 1.

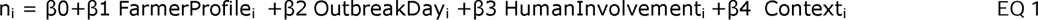

In this model:

- FarmerProfile is a categorical variable that characterizes different farmer types, influencing their likelihood of taking action based on predefined archetypes.
- OutbreakDay is a continuous variable denoting the day of the outbreak, which may reflect time-dependent factors influencing action.
- HumanInvolvement is a binary variable indicating whether human involvement in the outbreak is present.
- Context is a binary variable representing whether the context of the outbreak is favourable for action.

From the fitted beta regression model, we generated predictions for the daily probability of action using the model’s parameter estimates and the observed values of the predictor variables. Predictions were computed on the scale of the response variable (probabilities between 0 and 1) by applying the inverse logit transformation to the linear predictor (n_i_). These predictions represent the expected probability of action for each day, conditioned on the specified context, human involvement, outbreak day, and farmer archetype. The resulting daily probabilities were visualized over time to highlight trends and variations in the likelihood of action, allowing us to identify patterns or influential predictors in outbreak dynamics. All computations and visualizations were performed in R (version 4.3.2) [25], and functions from the *betareg* [26, 27] and *ggplot2* [28] packages.

### Estimation of probability of transmission/spread: Disease transmission model

To estimate the daily probability of disease transmission to another farm, we developed a two-tiered model to simulate disease dynamics within and between pig farms (see Supplement S2). The within-farm model used a stochastic Susceptible-Infectious-Recovered-Maternally immune (SIRM) compartmental model implemented with the R package *SimInf* [29]. It incorporated population and disease dynamics, such as birth rates, age group transitions, and disease progression. Parameters were based on literature [30–32], and variability in outcomes (e.g., outbreak extinction vs. endemicity) was captured through stochastic simulations. The median infectious duration from farms with 200 sows informed the between-farm model. The within-farm model also informed the on-farm outbreak trajectory that was presented to the stakeholders to measure the probability of action.

The between-farm transmission dynamics were captured by simulations of a distance-based transmission kernel model [33], where infection probability depended on inter-farm distance and infectious duration. Using farm location data from the Netherlands, the model was parameterized via Approximate Bayesian Computation to match an end-prevalence of 40% [34]. Sensitivity analyses tested the effects of varying key parameters. This integrated framework provided insights into transmission dynamics at both within-farm and between-farm scale.

The probability of transmission from an infected farm was considered the relevant (unfavourable) outcome and was based on the transmission kernel described above (and in Supplement S2 for details). We considered the daily probability of transmission to another farm. Since this probability depends on the density of farms around an infected farm, we randomly sampled two farms that were representative for a Dutch setting: One in a high dense area with more than 50 farms with a 5 km radius, and one in a low dense area with less than 25 farms (See supplement Figure S5). The transmission probability to any other farm was calculated based on the fitted transmission kernel, after which the product of the complement probability was calculated. One minus this probability was interpreted as the probability that at least one other farm got infected during that day. Within these two settings we additionally considered a scenario where the daily probability of transmission was uniform over time, and one where the cumulative probability after 100 days was similar to the uniform distribution, but the daily risk was scaled to the daily outbreak size within the farm as a proportion of infectious animals. Thus, for the daily probability of between farm transmission we ended up with four scenarios: A ‘high dense uniform’, ‘high dense proportional (to outbreak)’, ‘low dense uniform’ and ‘low dense proportional’ scenario.

### Competing probabilities: Between farm transmission versus detection

To assess whether there was disease detection before spread to other farms per scenario, we considered two competing probabilities: The probability that the infection spreads to another farm (pspread) and the probability of detection (p_detection_). The latter is the result of p_action2_ (probability of notification of the veterinarian) which is conditional on p_action1_ (probability of noticing symptoms) or p_detection_ = P(Action2|Action1). We thus assumed that successful detection was achieved when both actions were taken. For each combination of scenarios, we ran 1000 simulations in which we sampled for 100 days from the daily probabilities, where we assumed a Bernoulli trial with a probability p. The first day for which we sampled a ‘success’ was considered that the day the event (detection or spread) took place. We then compared the day at which spread occurred with the day that the detection occurred. When spread occurred before detection, we considered this an ‘outbreak’. We report the percentage of simulations that resulted in outbreaks per scenario. In total we compared 48 scenarios (Table 3).

**Table 3.** Overview of the scenarios for the probability of detection (pdetection) and the probability of between farm transmission (p_spread_).

| Outcome | Variable | Levels (n) |
| --- | --- | --- |
| P_detection | Farmer profile | Three farmer archetypes (3) |
| P_detection | Human involvement | Yes/no (2) |
| P_detection | Context | No information/Outbreak across the border (2) |
| P_spread | Farm density | Low/high (2) |
| P_spread | Daily risk profile | Uniform/proportional to on farm infections (2) |

## Results

### Probability of action

The control questions from the survey showed overall consistency, with participants largely aligning on the ‘Family-oriented farmer’s’ external influence, resistance to adopting new technologies, and confidence in established methods. However, divergences were noted in perceptions of subjective norms for the ‘Business oriented farmer’ and the ‘Farmer without successor’ profiles, likely due to differing interpretations of the influence by others. The full interpretation of the control questions is provided in the Supplement S3.

We found that the estimated median probability of noticing symptoms on day 10 of the outbreak was 0.6 (range: 0.25-0.9), increasing to 1 (range: 0.93-1) by day 20. The estimated median probability of reporting disease to the veterinarian was 0.6 (range: 0.25-0.85) on day 10 and 1 on day 20 (range: 0.87-1). Part of the variability between the estimates within the same day was caused by the different scenarios, and the different farmer profiles. However, between-respondent variability was present as well (Figure 3).

**Figure 3.**
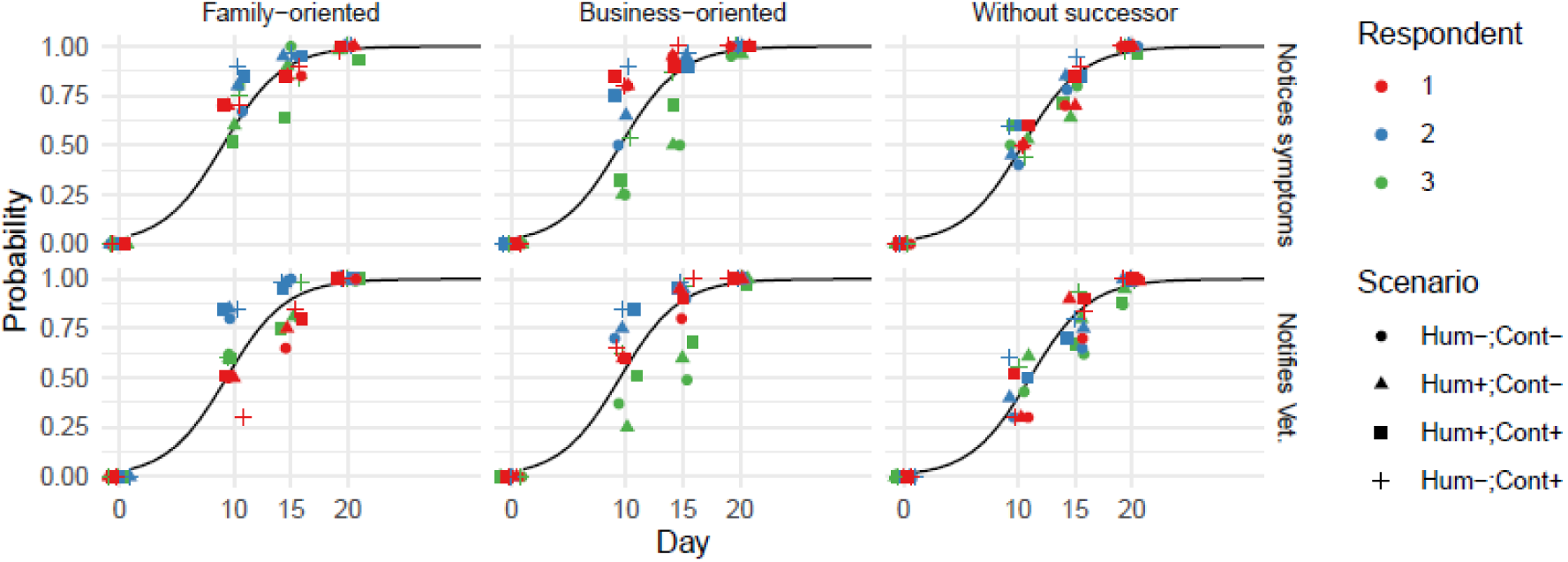
Probability of action as function of time in the outbreak. The smooth lines show the predicted probability from the fitted beta-regression model, the points show the response for each scenario, and time-point, stratified by respondent (colour) and scenario (shape). Panels show the farmer profile (columns) and action of interest (rows). Abbreviations: Hum: Human involvement in the outbreak: Yes (+) or no (-), Cont : Contextual information about earlier outbreaks known (+) or unknown (-).

The beta-regression showed a strong relation between the extent of the outbreak and the probability of both actions: For each day later in the outbreak, the expected log-odds increased by 0.38 for noticing symptoms, and for notifying the veterinarian (Table 4). Similarly, the farmer profile ‘No successor’ showed a lower likelihood for both actions. Human involvement and context were judged to be drivers of the probability of actions for some respondents (Supplement 3, individual results), but overall they did not significantly change the probability of action. From this followed a median time to notification of the veterinarian after an on-farm introduction of an infection of 8 days (95% CI: 4-12 days) for the ‘Family-oriented farmer’, 9 days (95% CI: 5-12) for the ‘Business-oriented farmer’, and 10 days (95% CI: 6-13) for the ‘Farmer without successor’.

**Table 4.** Effects of farmer profile, time in the outbreak (Day), human involvement and context knowledge on the probability of noticing symptoms and the probability of notifying the veterinarian as obtained by beta-regression.

| Variable | Estimate | Std. Error | Z-value | P-value |
| --- | --- | --- | --- | --- |
| **Outcome: Noticing symptoms** |  |  |  |  |
| (Intercept)* | -3.467 | 0.182 | -19.093 | 0 |
| Farmer: Business-oriented | -0.179 | 0.173 | -1.033 | 0.301 |
| Farmer: Without successor | -0.520 | 0.180 | -2.895 | 0.004 |
| Day | 0.383 | 0.016 | 23.772 | 0 |
| Human: TRUE | -0.193 | 0.142 | -1.362 | 0.173 |
| Context: TRUE | 0.185 | 0.143 | 1.299 | 0.194 |
| **Outcome: Notifying the veterinarian** |  |  |  |  |
| (Intercept)* | -3.692 | 0.217 | -17.021 | 0 |
| Farmer: Business-oriented | -0.062 | 0.168 | -0.368 | 0.713 |
| Farmer: Without successor | -0.658 | 0.177 | -3.717 | 0.0002 |
| Day | 0.386 | 0.016 | 24.643 | 0 |
| Human: TRUE | -0.081 | 0.140 | -0.580 | 0.562 |
| Context: TRUE | 0.235 | 0.141 | 1.670 | 0.095 |

\*The profile for the 'Family-oriented farmer' served as reference category.

### Probability of between farm transmission

We found that the daily probability of transmission to at least one other farm in a high-dense area was approximately a 4-fold higher than in a low-dense area (4.7e-3 vs 1.2e-3, Figure 4). Given these probabilities, 5% of the farms in a similar setting would have caused an infection in another farm after 11-49 days, depending on the scenario (Table 5).

**Figure 4.**
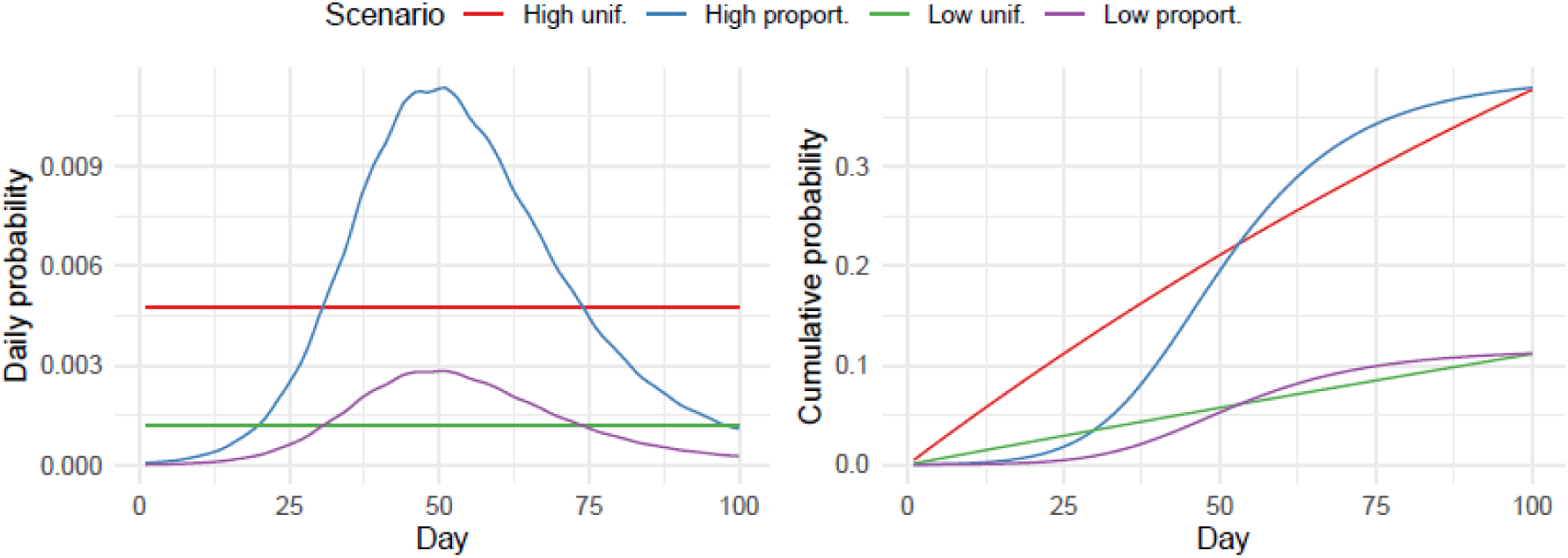
Probability of farm-to-farm transmission over time. The daily (A) or cumulative (B) probability that an infected farm infects at least one other farm over time. The scenarios are based on the farm- density (high vs low) and assumed distribution of the risk over time (uniform [unif.], or proportional [proport.] to the number of infectious animals on the farm).

**Table 5.** The number of days after the introduction of the infection on the farm for which 1, 5, or 10% of the cumulative risk of spill-over to at least one other farm was reached by scenario based on the farm- density (high vs low) and assumed distribution of the risk over time (uniform, or proportional to the number of infectious animals on the farm).

| Scenario | Days at 1% risk | Days at 5% risk | Days at 10% risk |
| --- | --- | --- | --- |
| High uniform | 2 | 11 | 22 |
| High proportional | 21 | 33 | 40 |
| Low uniform | 9 | 43 | 89 |
| Low proportional | 31 | 49 | 76 |

### Conditions under which successful early detection was achieved

We found that the proportion of on-farm outbreaks of zoonotic swine influenza that resulted in spread to other farms before the veterinarian was notified, was the highest in the pig-high density areas, where the risk was assumed to be uniformly distributed (Figure 5). There 3.6-4.2% of the on-farm outbreaks resulted in transmission to at least one other farm before the veterinarian was notified. The behavioural drivers associated with the different farmer profiles did result in nuanced differences in outbreak risk. The profile of the ‘Farmer without successor’ returned the highest outbreak risk.

**Figure 5.**
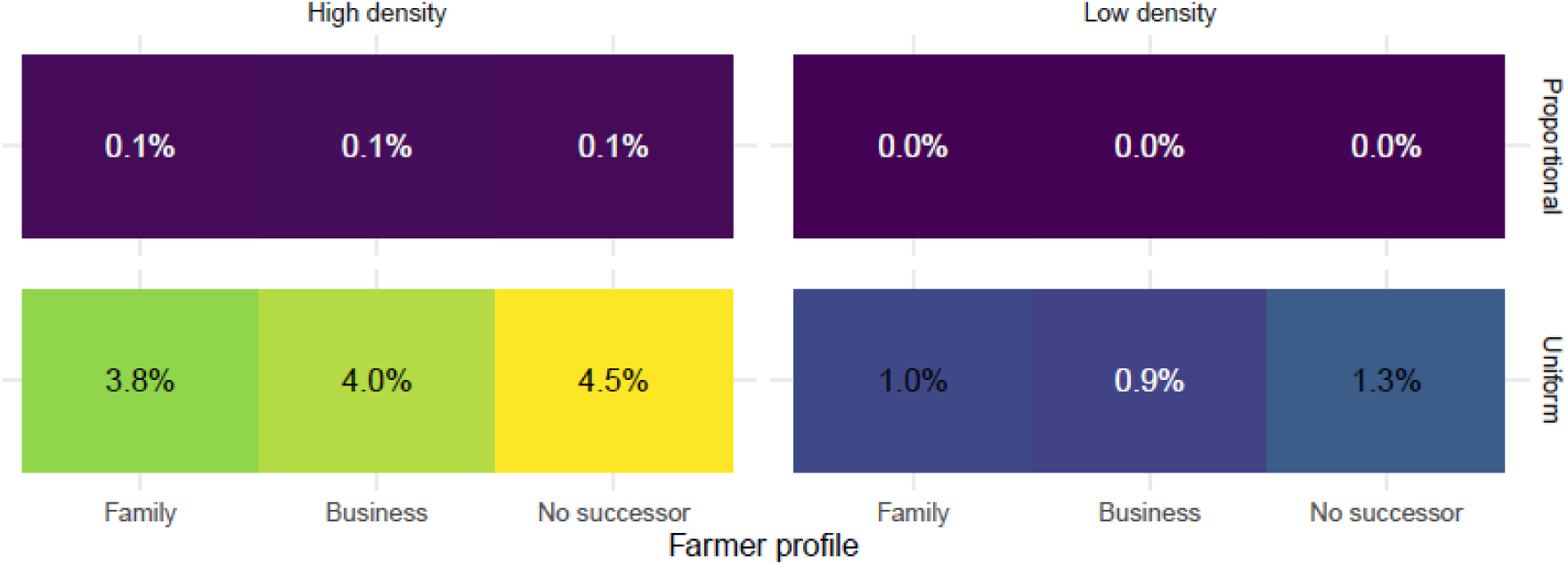
Proportion of outbreaks that spread to other farms before detection stratified by Farmer profile. Facets: Between farm risk (columns: high density vs low density areas; rows: proportional with number of infectious animals, or uniformly distributed risk). X-axis: Farmer profile (Family oriented [Family], business-oriented [Business], without successor [No successor]).

## Discussion

Here we have shown that we can successfully integrate concepts from behavioural science, disease transmission modelling and epidemiology to create more nuanced estimates of the effectiveness of interventions in the pandemic preparedness domain. We described a method that allows to first estimate probabilities of transmission and the required human actions that would counter transmission. We then sampled from these competing probabilities to arrive at a single metric of the probability of an outbreak. For the use-case ‘early detection of zoonotic swine influenza’, we found that it was estimated that 95% of the farmers would notify their veterinarian within 13 days after an infection was introduced on a farm. Both the probability that farmers notice symptoms and report disease to the veterinarian were driven by the extent of spread, expressed as the number of days after the infection was introduced on the farm. The speed with which farmers were thought to act also depended on their profile, where the ‘Farmer without successor’ was less likely to act. Human infections linked to the outbreaks were not significantly altering the proportion that took action. Farm density, but mainly the assumption on the shape of the distribution of the probability of transmission to another farm, drove the proportion of outbreaks that spread to one or more farms before the veterinarian was notified.

The results of the case-study indicate that increasing the understanding of the transmission and response in high risk areas is crucial to be able to perform targeted (risk-based) interventions, which is in line with earlier findings [35, 36]. Predicting how people respond to interventions in these settings, based on theories embedded in the social sciences will help identify weak spots, and will result in an even finer scale of targeting of interventions. Communication style and the implementation of the interventions can be aligned with the needs of the target groups [14].

A main strength of the framework that we present here, is that it follows a generic line of thought, making it applicable to any setting where human behaviour is required to counter disease transmission. Through-out the field of outbreak and pandemic preparedness, many of the interventions require people to take certain actions, whether it is people opting to vaccinate, notify or test for disease, or adjusting their behaviour to diminish disease transmission risk. With the framework we make each step explicit and thus transparent, helping policymakers come to decisions based on best available evidence. For researchers and practitioners, it helps identify the sources of greatest uncertainty, and thus where research or additional data collection is needed.

Our work has several limitations as well. What we present here as a probability of action, is a behavioural intention as judged by experts. The disconnect between intention and actual behaviour is a well described phenomenon [37]. However, in a hypothetical setting of preparedness for an expression of a disease that has not been encountered yet, it is hard to quantify this gap, simply because the behaviour has not been observed yet. Historic data on the outbreak response of earlier outbreaks might offer some insights, however differences in context will always remain.

Choosing to work with farmer profiles was well-adapted to the study and available resources; however, this approach presents several limitations to be acknowledged. Firstly, these profiles can oversimplify the diversity of behaviours and decision-making processes among farmers, potentially leading to generalized conclusions that do not reflect individual circumstances. Additionally, they do not account for cultural and regional variability, which are critical factors influencing farming practices. To address these limitations in future studies, qualitative research methods such as interviews or focus groups could be employed to gather deeper insights into the motivations and experiences of various farmer groups.

To tackle the limitation of the indirectness caused by relying on expert judgement, we can replace this by a well-designed survey, where stakeholders are targeted directly. There we could directly measure the behavioural drivers [38–40] and their response to the different outbreak scenarios. Assuming that one would be able to recruit a sufficient sample size, stakeholders can be asked on which day they judge that they will take a certain action. The resulting ‘time-to-action’ data that is thus obtained can then be analysed using survival analysis. We must be aware that there remains a risk of anchoring when performing expert/stakeholder elicitation [41], where the order of the scenarios that are presented to the stakeholder might thus influence the results. Randomization of the order in which scenarios are presented can address this. Exploring how more immersive methods, such as serious gaming [42] or virtual reality [43], of presenting outbreak scenarios could trigger a more ‘life-like’ response would be a logical extension of the work.

Here, we presented and tested a framework for integrating concepts from behavioural sciences and epidemiology in which we can consider the full chain of actions required to achieve a desired outcome. We have arrived at a level of detail of model integration which is pragmatic and applicable, as illustrated by the use-case. Applying this methodology will support decision-makers in their efforts to develop evidence-based policy for outbreak and pandemic preparedness in which they can consider evidence from both social sciences and epidemiology.

## Acknowledgements

We would like to extend our gratitude to the veterinary swine specialists Dr. Lucia Dieste Pérez, Dr. Tijs Tobias of Royal GD, and Dr. Tosca Ploegaert of Wageningen Bioveterinary Research, and to Dr. Linda Peeters for piloting earlier versions of the survey instrument. We thank Gert Jan Boender of Wageningen Bioveterinary Research for providing expert input on the kernel-based approach of the between-farm transmission.

## Supplementary material

- S1. Survey
- S2. Model description
- S3. Detailed results/interpretation

## S1. Survey

### Introduction survey

To better understand and predict how a disease outbreak is detected early, understanding human behaviour is essential. We (researchers from Wageningen University and Research) are working on a methodology to measure behavioural intention (the intention to act) and link it to disease spread models. This way, we can better prepare for outbreaks of (animal) diseases.

The purpose of this questionnaire is to test the methodology to estimate the relationship between stakeholders’ willingness to act under different circumstances. In this questionnaire, we present various scenarios. We describe a type of personality (character), a type of outbreak, and contextual information.

We ask you to estimate the willingness to act of a character who is experiencing a certain disease outbreak. We ask you to put yourself in this persona’s shoes and, using the provided information, make an estimate at three different points in time during the outbreak. We are interested in whether you think it is likely that the character will recognize the disease (action 1) and subsequently report the disease to their veterinarian (action 2).

The described persona and scenarios are hypothetical and are solely for testing the methodology. The results will be analysed anonymously, with the respondent being identified only by a random, non-traceable identification number.

The survey consists of four parts: 1) this introduction, 2) the introduction of the persona, the description of the disease outbreak and the context, 3) control questions about the persona, 4) four different scenarios.

### Introduction persona, disease outbreak and context

**Persona (Farmer profiles):**

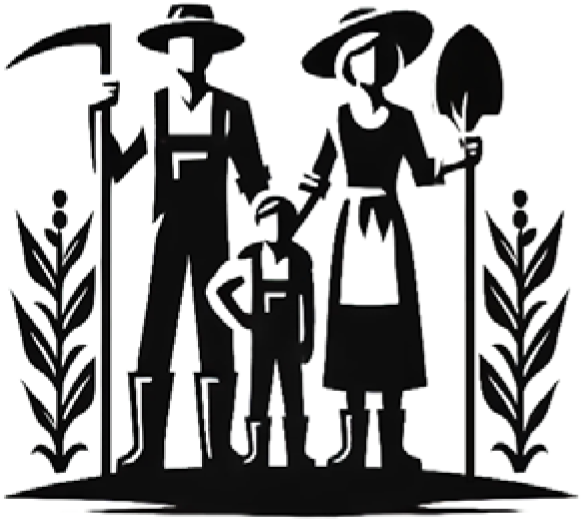

**A:** The **family-oriented farmer** starts work at the pig farm before dawn. He is committed to **preserving agricultural heritage** through technical skills and **generational knowledge**, while prioritizing **environmentally sustainable practices.** The large farm, with **significant sales**, is a family-driven operation focused on traditional farming, avoiding diversification and activities like agritourism.

Fellow farmers, neighbours, and family members’ opinions play a **significant role** in shaping their practices and decisions.

They feel **confident** managing the farm efficiently using **proven techniques**, which gives them a **sense of control** .

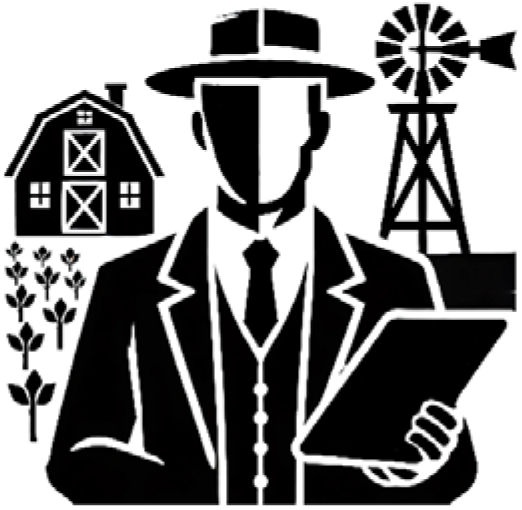

**B:** <u>The</u> **<u>business-oriented farmer</u>** focuses on **optimizing production** and **maximizing profits** . Running a large, efficient farm connected to global markets, they invest in modern housing, automation and equipment. External labour, both domestic and from EU member states, is crucial for operations.

**Market demands, industry trends,** and **financial stakeholders** shape their decisions. They maintain **strong networks** with other successful farmers and agribusiness experts, relying on industry and financial advice for their strategies.

This farmer focuses on **maximizing output** and **cutting costs** through **advanced technology** and **data-driven strategies** . Confident in managing the farm with **high-efficiency systems**, they view external labour as a key input factor and are less concerned about obstacles like **getting bank loans approved**, or **encountering issues with insufficient knowledge and available resources** .

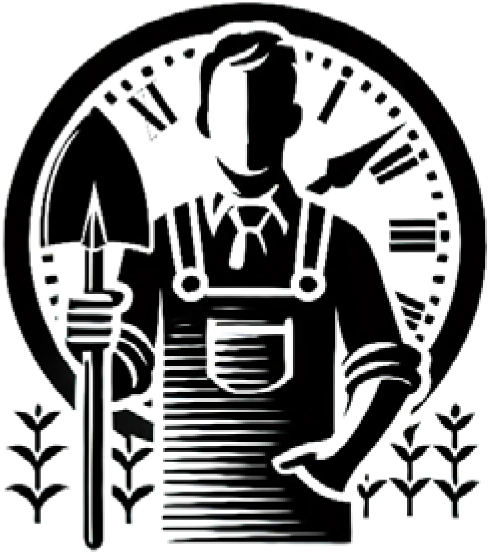

**C: The fa<u>rmer without successor</u>** focuses on **avoiding expenditures**, prioritizing **immediate profit over long** -**term sustainability** or ethical concerns. With **no foreseen successor**, their actions are driven by practical, **short** -**term goals** .

This farmer is influenced by **others who also value efficiency** and financial results. He is **less concerned** with the **views of those who emphasize sustainability** or **ethical practices** .

The farmer feels **overwhelmed** by **farming’s challenges** and **uncertainties**, lacking confidence in making decisions due to **limited resources** and **weak support networks**. This mindset leads to **hesitation**in **adopting new practices** or **technologies** .

**Disease outbreak:** The pig farmer works on a closed pig farm. It is a farm with 300 sows. At some point, with the purchase of new gilts, a disease is introduced to the farm. Increasingly more animals become sick: they have fever, they cough, and their appetite is reduced. It starts with a few animals coughing, but soon a larger outbreak develops (see table below).

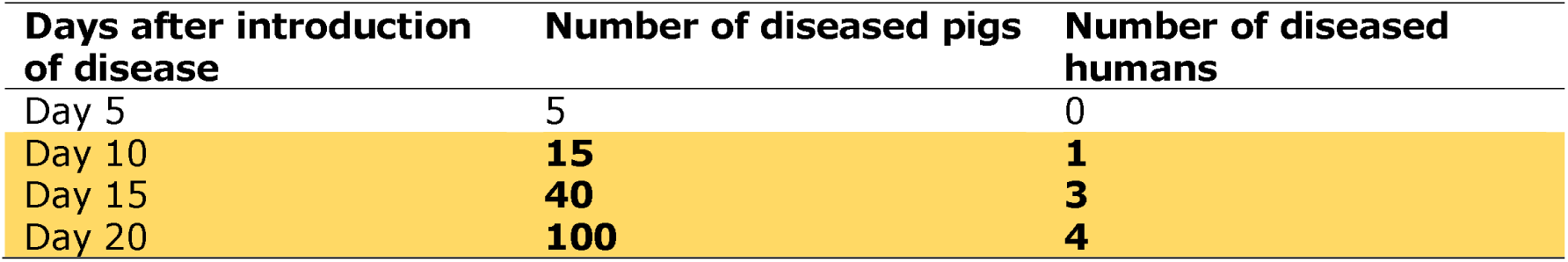

**Human involvement in the outbreak:** We consider two possibilities:

- **A:** Only pigs get sick.
- **B:** As time progresses, people also become sick, initially an employee, but later also people with whom the employee has been in contact. The pig farmer is informed about this (see table above, column ‘ Number of diseased humans’).

**Context:** The pig farmer is aware of the current situation regarding animal diseases. We consider two possibilities:

- **A:** There is a ’normal’ threat of animal disease outbreaks, there are no concerning outbreaks near the pig farm.
- **B:** There is an increased threat: Just across the border, there has been an outbreak of a ’flu-like’ swine disease that has also affected humans.

### Outcome

We are interested in two actions necessary to detect the disease outbreak: 1)The pig farmer notices that there is a disease of interest is spreading that may need to be reported (’notices’). 2) The pig farmer contacts his or her veterinarian (’reports to vet’).

We ask you to answer how likely you think it is that the pig farmer will notice the disease and subsequently reports the disease to their veterinarian. We use a scale to express the probability:

In part 3 of the survey, we ask some control questions about the persona. In part 4, we present a different combination of persona, outbreak, and context each time and ask you to estimate the likelihood of the outcome.

### Control questions

You will be presented with three questions to assess your understanding of the farmer profile.

[Control question relating to attitude]: Please rate your agreement with the following statement: “The farmer uses new technologies”:

- Strongly disagree

- Disagree

- Somewhat disagree

- Neutral

- Somewhat agree

- Agree

- Strongly agree

[Control question relating to subjective norms] Please rate your agreement with the following statement: “The farmer’s decisions about farm management are influenced by the opinions and expectations of others”:

- Strongly disagree

- Disagree

- Somewhat disagree

- Neutral

- Somewhat agree

- Agree

- Strongly agree

[Control question related to Relates to perceived behavioural control] Please rate your agreement with the following statement: “The farmer has high confidence in their farming methods”:

- Strongly disagree

- Disagree

- Somewhat disagree

- Neutral

- Somewhat agree

- Agree

- Strongly agree

### S2. Disease transmission model

Infectious disease dynamics within a pig farm are governed by different processes. Both the disease properties and the farm management influence the natural course of a disease outbreak. Farm management, for example, drives the intensity of contact between animals and the frequency with which new susceptible individuals are added. Within the farm, we modelled the disease dynamics using a stochastic compartmental model. Between pigs farms we considered that the disease dynamics could be described using a transmission kernel [33]. We constructed both a ‘within farm model’ and a ‘between farm model’, the infectious duration, the time that at least one infectious animal was present on the farm, was used to fit the between farm model; the between farm model informed the probability of transmission to another farm at farm level (Figure S2.1).

### Within farm model

We use a compartmental model in which dynamics between different disease states are expressed by ordinary differential equations. Here, animals can exist as susceptible, infectious, recovered or maternally immune individuals (SIRM), similar to [30, 31]. Population dynamics, birth, ageing, and moving out of the farm are implemented as ’post-time-step events’ as defined in the R package SimInf [29]. See Figure S2.2 for a graphical depiction of the model. In the baseline model, we assume homogeneous mixing between groups and transmission (β) and recovery (γ) rates to be the same for all groups. In order to calculate the disease incidence (or new cases per age group), we introduce a dummy compartment (C, not depicted in the figure) which represent the cumulative incidence.

Population dynamics were implemented as SimInf ‘events’, in which groups of animals move to a subsequent age-group at discrete timepoints. We assumed 2.4 litters per sow per year, with an average litter size of 14 piglets. Piglets were weaned after 30 days, and after another 40 days they moved to the fattening group. Sows were replaced with susceptible new sows with a rate of 0.4/year (table S2.1).

Parametrisation of the model was based on literature existing literature (Table S2.1). Disease progression was modelled based on the number of incident cases (C). After a latent period of 2 days, animals remained symptomatic for 5 days. We allowed 175d as burn-in period, to populate the farm based on an initial number of sows. After the burn-in period, 5 infectious sows were introduced. Figure S2.3 gives an example of the median number of symptomatic animals from the day of introduction for a closed farm of 200 sows.

Due to the stochastic nature of the disease, we observed that a proportion of the introductions resulted in outbreaks that died out (Figure S2.4), and a proportion in endemic circulation within a farm, where the virus continued to circulate due to population renewal and loss of (maternal) immunity. These proportions were depended on the farm size, in line with earlier observations [31]. The simulated infectious duration of a farm with 200 sows, or a total farm size of 2795 pigs, given a closed farm, was used to estimate the parameters for the between farm model.

### Between farm model

To model the transmission between farms, we applied a kernel based approach [33]. Transmission is assumed to decrease with the distance between farms and the transmission kernel describes the daily transmission hazard (h) between an infectious and susceptible farm as a function of, the Euclidean distance between an infectious (i) and a susceptible farm (j), r0 is the ‘kernel offset’, h0 is the amplitude of the transmission kernel indicating the transmission hazard for very small distance and is a scaling exponent (EQ S2.1).

(EQ S2.1)

The probability that an infected farm infects a susceptible farm is a result of the infection hazard (h) and the infectious duration of the infected farm (T_i_) (EQ S2.2) [45].

(EQ S2.2)

From that follows that the probability that a single infected farm will infect at least one other farm within a certain time (Ti) is given in Equation S2.3, or one minus the probability that all farms escape infection.

(EQ S2.3)

To parameterize the model, the kernel was implemented in a stochastic framework in which we considered multiple stochastic events: Transmission or escape from infection, and in case of infection: recovery or escape from recovery. In case of infection, the infectious duration was sampled from distribution of the infectious duration of the within farm model for a farm of 200 sows.

We used data location data of pig holdings in the Netherlands on which more than 100 pigs were present (Figure S2.5). We assumed (1) no distinction between farm types, (2) no cross immunity for the emerging zoonotic strain, (3) no within-host competition between emerging and existing strains, and (4) infected farms are immediately infectious (no latent period at farm level).

The kernel was fitted to an end prevalence where 40% of the Dutch farms experienced infection [34], with an assumed r0 (the ‘kernel offset’) of 1, and a scaling exponent () of 1.6, based on earlier experience [33]. Infection was seeded in a low pig-dense area and a high pig-dense area (Figure S2.5). An Approximate Bayesian Computation (ABC) approach was used to parameterize the kernel amplitude (h0). We evaluated the effect of the kernel offset (default of r0=1) on the proportion of introductions that lead to endemic circulation and the end-prevalence given endemic circulation, by increasing and decreasing the value to 1.5 (higher r0) and 0.5 (lower r0) (Table S2.2, Figure S2.6). The values were robust against these changes?.

### S3. Detailed results: Response per participant

#### ‘Validation’ of behaviour survey

Figure S3.1 presents the responses to three validation questions directed at survey participants regarding the different farmer profiles. The questions address three constructs of the TPB: (1) subjective norms, exploring whether the farmer’s management decisions are influenced by others’ opinions; (2) attitudes towards innovation, assessing the likelihood of adopting new farming technologies; and (3) perceived behaviour control, gauging the farmer’s confidence in their own farming abilities. These questions serve to verify if respondents had a consistent understanding of the farmer profiles.

The responses suggest that participants broadly shared similar perceptions of the profiles. For instance, all respondents agree that the “family” farmer’s decisions are influenced by external opinions (e.g., family, friends), that this farmer is unlikely to adopt new technology, and that they have confidence in their established methods. Across seven of the nine instances, responses fell consistently along a similar spectrum of agreement or disagreement, indicating an aligned understanding of the behavioural traits associated with the profiles.

However, divergence was observed in responses to the subjective norms question regarding the “entrepreneur” and “no successor” profiles. This inconsistency may stem from varying interpretations of “influence by others”. One hypothesis for these discrepancies could be that the concept of “influence by others” is interpreted differently depending on the respondent’s own perspective on entrepreneurial independence or social isolation in succession planning. The absence of precise guidance on what constitutes “influence by others” in the survey could have led respondents to rely on personal or cultural assumptions, adding to the variability in responses. The brevity of the profiles—kept short for survey efficiency—may also have contributed to these differences in interpretation.

For the “entrepreneur” profile, respondents may perceive entrepreneurs as inherently self-reliant. However, the profile description suggests a farmer influenced by financial stakeholders and a network of other successful farmers and agribusiness experts. This duality can create ambiguity, leading to conflicting views on the degree to which the entrepreneur farmer is influenced by others in their decision- making.

Similarly, the “no successor” profile may invite varied interpretations. The profile description suggests that the farmer is influenced by others who prioritise efficiency and financial outcomes, while paying less attention to those who advocate for sustainability or ethical practices. Some respondents might interpret a lack of succession planning as a sign of isolation or disengagement from community influences. Others, however, might interpret the lack of a successor as increasing a farmer’s reliance on external guidance (e.g., from advisers, neighbours, or community leaders) for day-to-day decisions, thus aligning with agreement on subjective norms.

This highlights the importance of defining concepts like “influence by others” more precisely in future surveys to minimise interpretative ambiguity.

---

**Figure S2.1.**
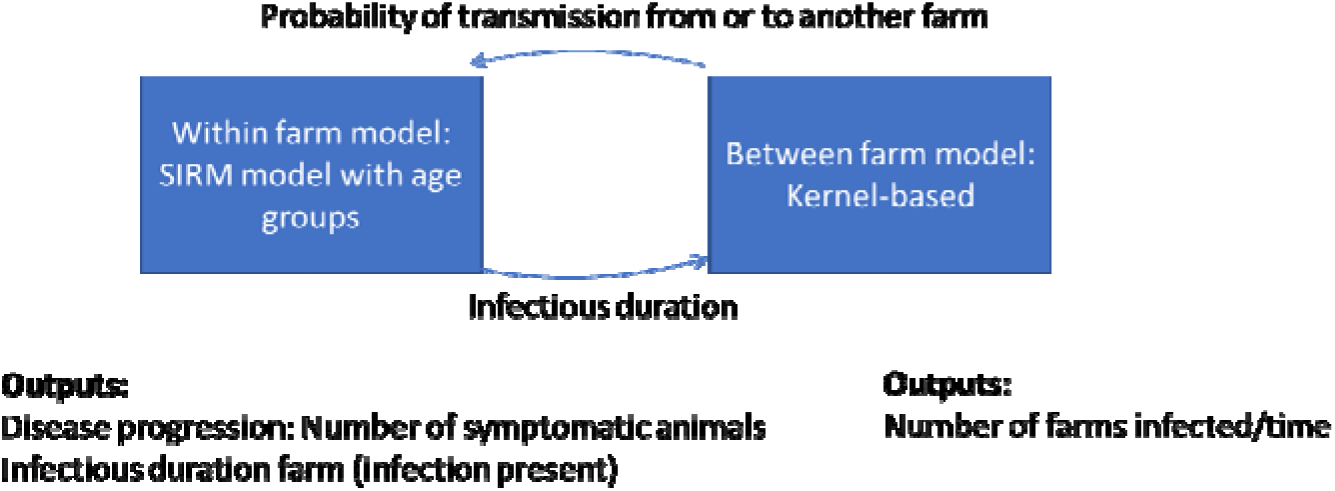
The interaction between the ‘within farm model’ and the ‘between farm model’. Output from the within farm model (the infectious duration) is used to parameterise the between farm model. Subsequently the between farm model informs the ‘probability of transmission to another farm’ at the farm level.

**Figure S2.2.**
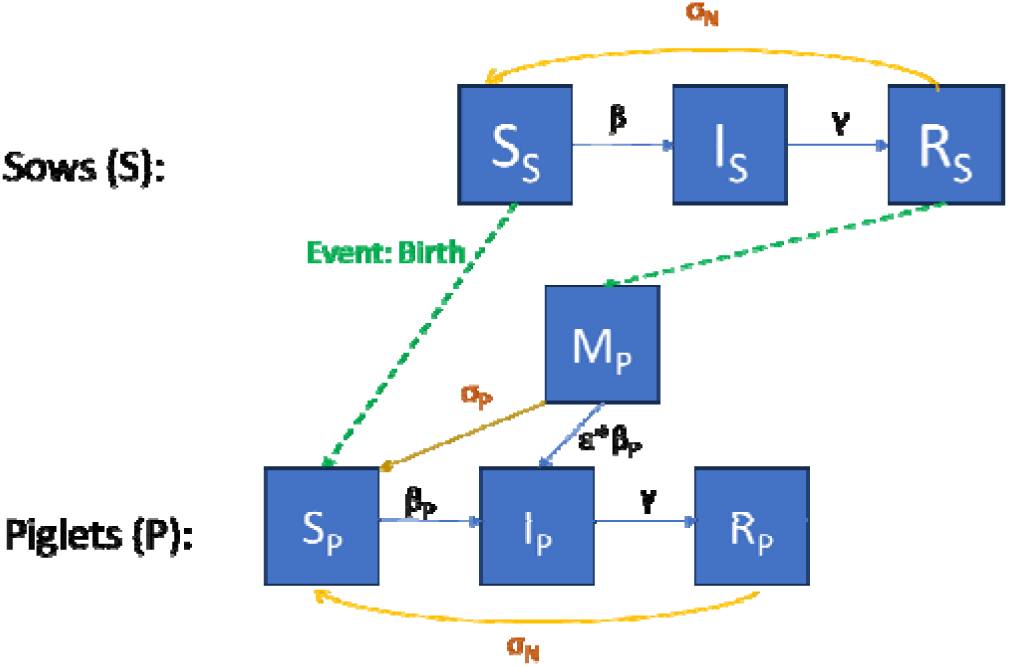
Graphical representation of the Susceptible-Infectious-Recovered-Maternal immune (SIRM) compartmental model in which animals exist in age-classes sow (S), piglet (P), weaned piglet (W, not shown here) and fattening pig (F, not shown here). Piglets born from immune sows (R_S_) have maternal antibodies (M_P_) that wane over a period of 11 weeks (σ_P_=1/77 days^-1^).

**Figure S2.3.**
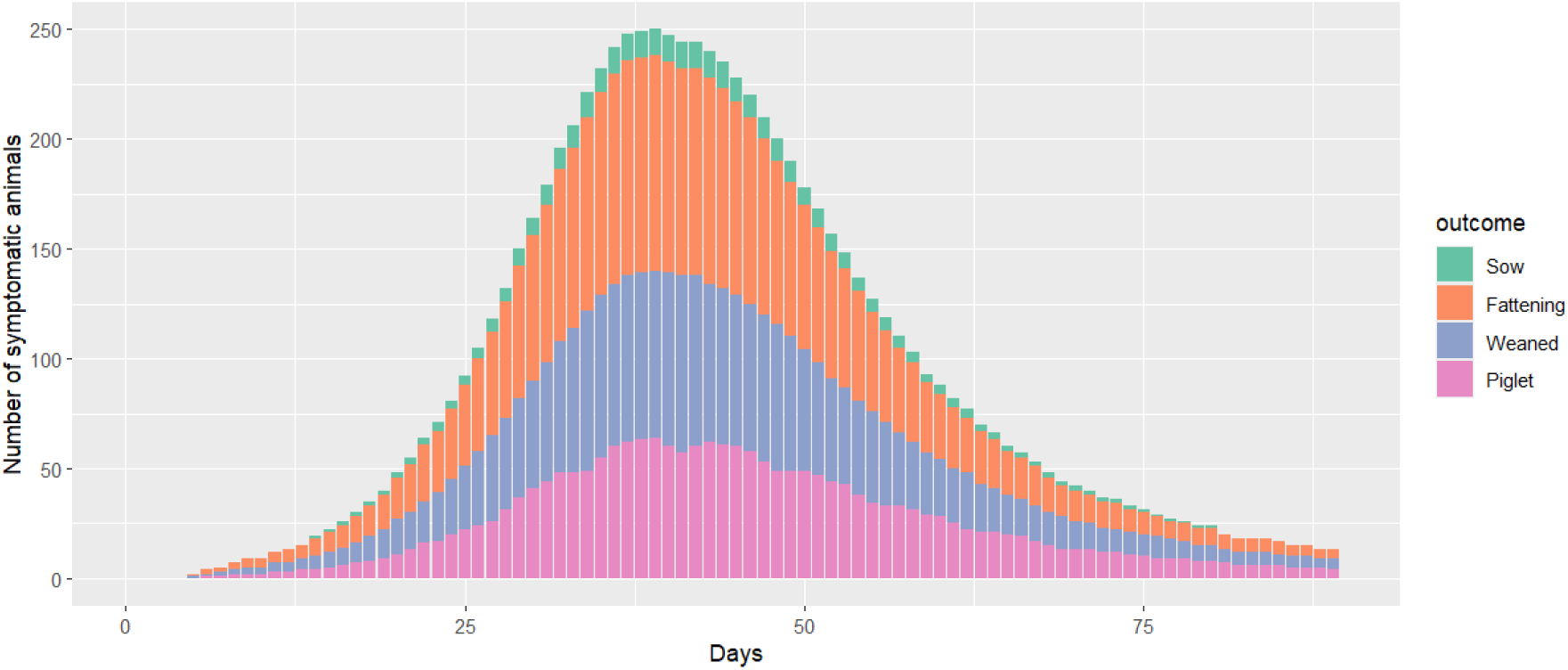
The median number of symptomatic animals per age group over time in a farm with 200 sows.

**Figure S2.4.**
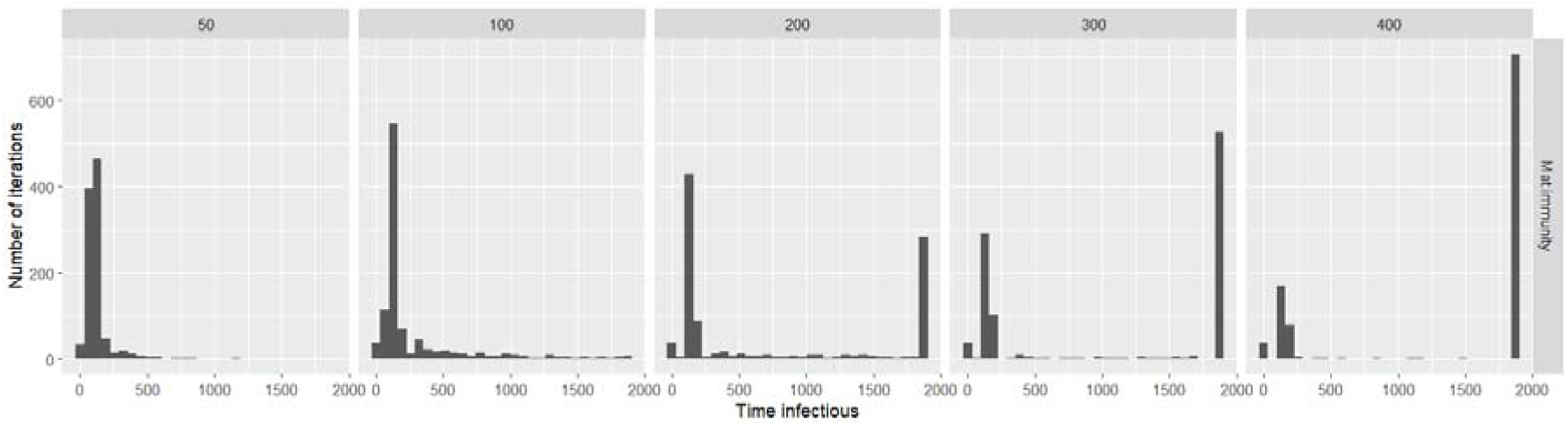
The duration of an outbreak (Time infectious) for 1000 iterations per scenario with number of sows per farm (50, 100, 200, 300, 400, as column facets).

**Figure S2.5.**
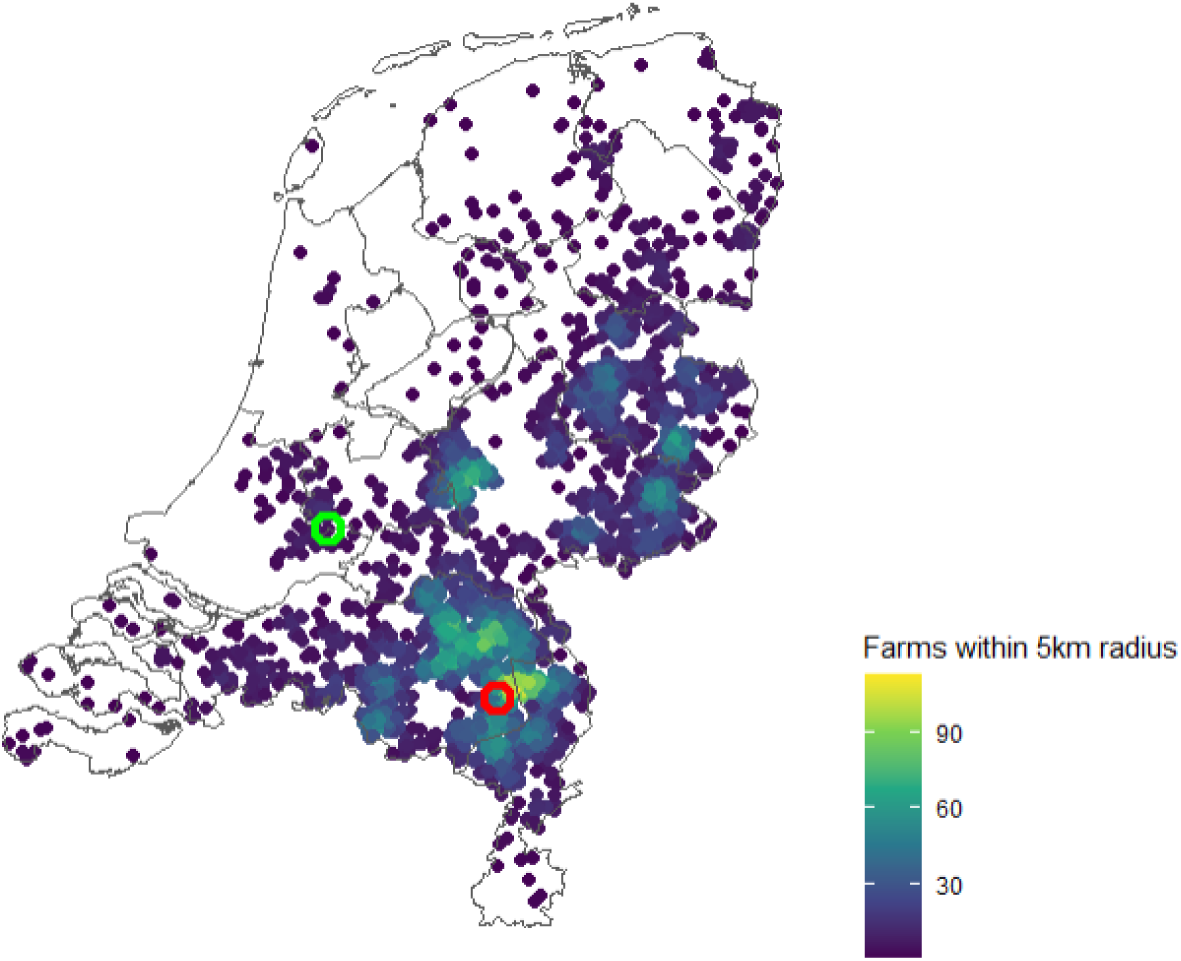
Distribution of pig farms within the Netherlands with more than 100 animals/holding (point) and the number of farms within a 5 km radius (colour), selected farms that represent a high- dense (red circle) and a low-dense (green circle) area.

**Figure S2.6.**
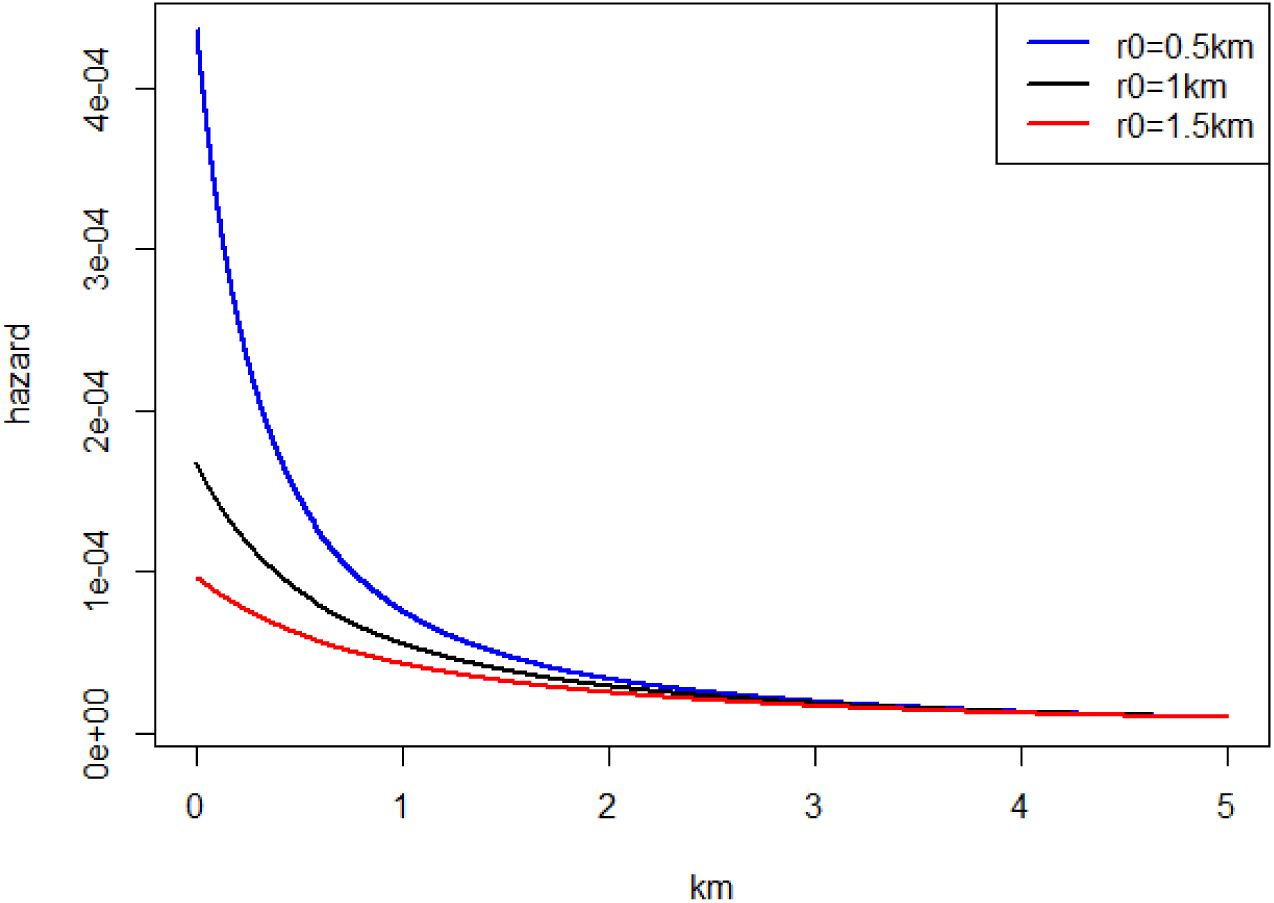
Visualization of the relationship between the infection hazard (y-axis) and distance (x- axis).

**Figure S3.1:**
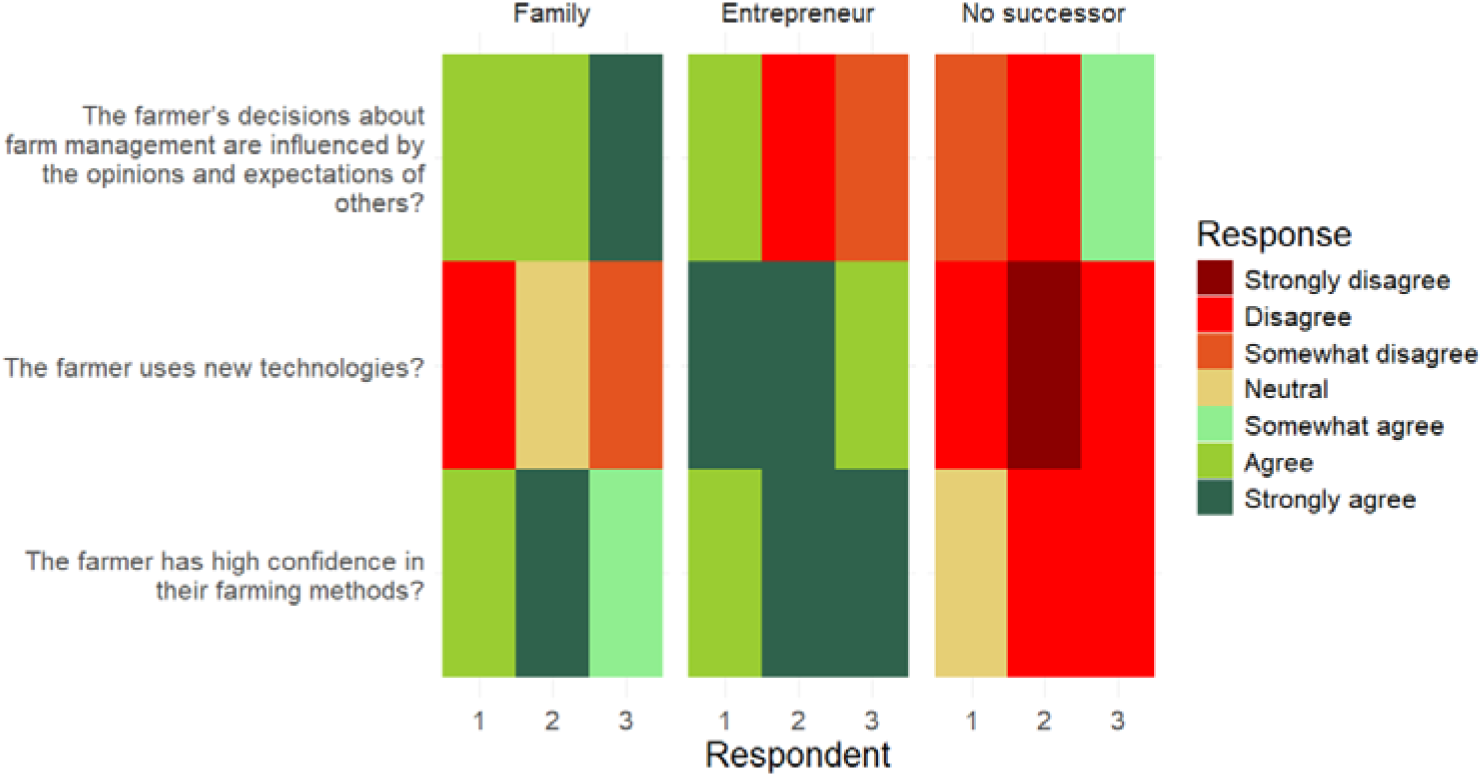
Survey respondent agreement on behavioural traits across farmer profiles.

**Table S2.1.**
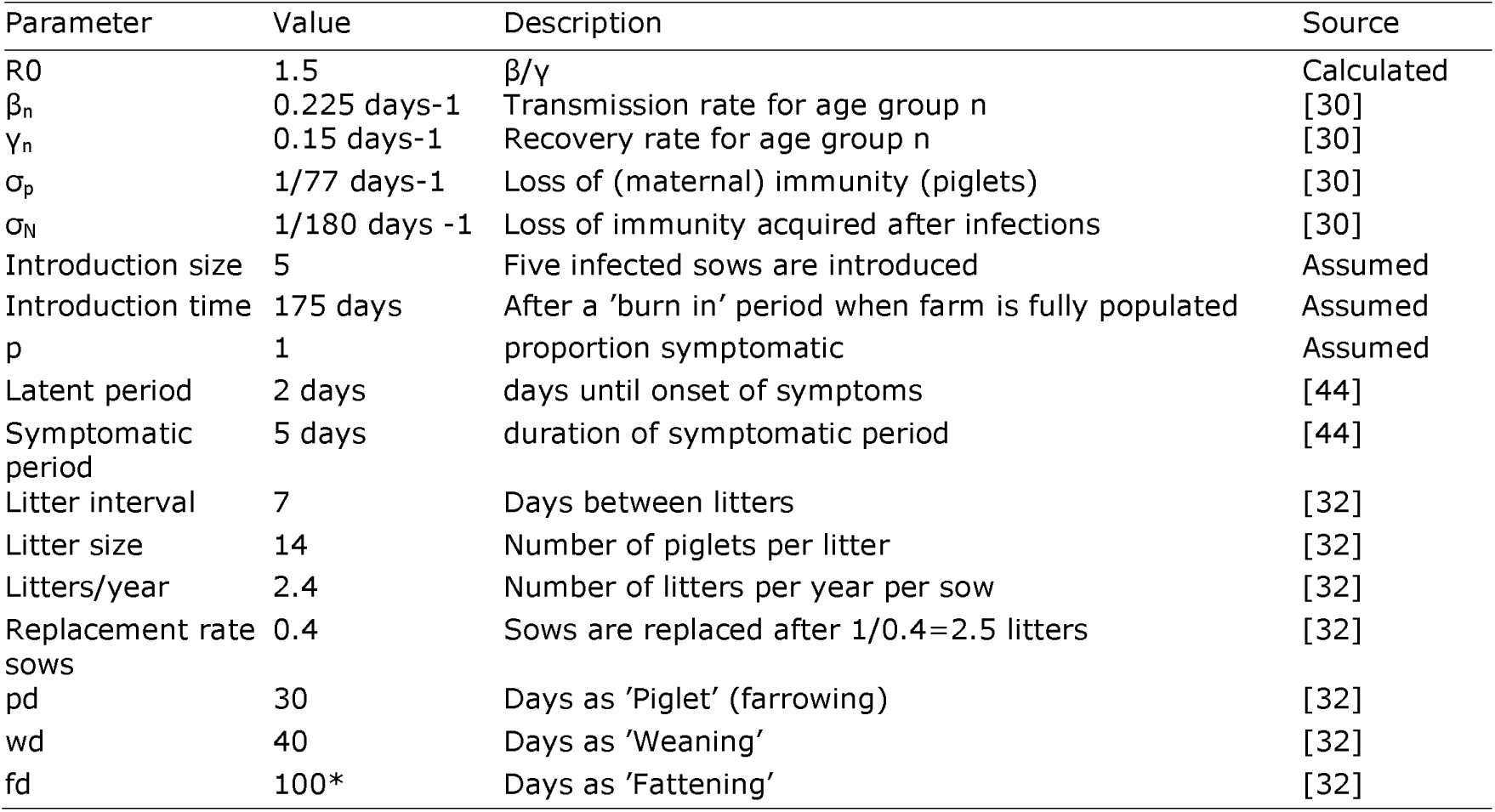
Parameters used to describe the disease dynamics and the population dynamics of the within farm model.

**Table S2.2.**
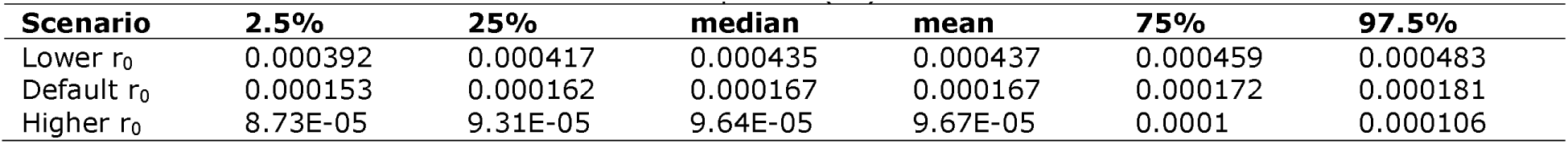
Parameter estimates for the kernel amplitude (h0).

